# Robust Estimation of Rotational Diffusion Tensors of Proteins from Molecular Dynamics Simulations

**DOI:** 10.1101/2025.05.27.656261

**Authors:** Simon L. Holtbrügge, Lars V. Schäfer

## Abstract

Rotational diffusion is a fundamental physical process that determines the rotational motion of proteins in solution. It plays a role, for example, in molecular association processes and in theories of spectroscopic experiments in solution. In addition to experimental methods, molecular dynamics (MD) simulations have emerged as a powerful method to investigate rotational diffusion. Diffusion models are fitted to rotational correlation functions extracted from the simulations. In this work, we conducted a time-dependent analysis of the model parameters prior to fitting, using extended all-atom MD simulations of ubiquitin as a model system. A comparison to Brownian dynamics (BD) simulations confirms whether the rotational dynamics observed in MD actually follow the theory of Brownian rigid-body diffusion. In addition, the analysis reveals correlation time intervals that are suitable for fitting anisotropic, semi-isotropic, or isotropic diffusion models. To this end, a two-step optimization scheme is employed that combines a global and a local search in parameter space. BD simulations are used to estimate uncertainties of the diffusion coefficients as well as directional uncertainties of the principal axes. We found that ubiquitin exhibits nearly semi-isotropic rotational dynamics, in good agreement with experimental NMR data. The approach is general and can be used to investigate, for example, the rotational diffusion of molecules in various biomolecular environments, or to compute NMR relaxation parameters of proteins. An implementation of the method is freely available at https://github.com/MolSimGroup/rotationaldiffusion.

## I. INTRODUCTION

Rotational diffusion models are frequently employed to characterize the rotational dynamics of molecules in the liquid phase, including soluble proteins,^1–5^ membrane proteins,^6–10^ or proteins in crowded environments.^11–19^ Furthermore, rotational diffusion models are used to describe the overall motion (“tumbling”) of molecules in the analysis of nuclear magnetic resonance (NMR) spin relaxation experiments.^20–23^ Rotational diffusion of proteins can be investigated using well established techniques such as NMR spectroscopy,^2,8,15–17,24^ fluorescence anisotropy spectroscopy,^5,18^ fluorescence correlation spectroscopy,^3,12,17^ or electron paramagnetic resonance spectroscopy.^9,11^ Molecular dynamics (MD) simulations complement laboratory experiments by providing atomistic resolution, which allows to study the coupling of rotational diffusion to other molecular processes, such as translational diffusion,^25^ or transient cluster formation.^13,14^

Studies on the rotational diffusion of molecules in solution commonly employ Favro’s diffusion theory, which is based on the Brownian diffusion of a free rigid rotor and utilizes a constant second-rank tensor to quantify rotational diffusion.^26,27^ The tensor can be expressed *via* its three principal diffusion coefficients >*D*_*x*_, *D*_*y*_, and *D*_*z*_, and their corresponding orthogonal principal axes. Diffusion is anisotropic if all coefficients are distinct, semi-isotropic if two coefficients are degenerate, and isotropic if all three coefficients are degenerate. If diffusion coefficients are degenerate, their principal axes are arbitrary (but orthogonal). Hence, the number of free model parameters decreases from six for the anisotropic case over four for the semi-isotropic model to one for the isotropic model.

To our knowledge, NMR is the only technique that has been successfully applied to extract the full anisotropic diffusion tensor of a protein from laboratory experiments, given that the protein structure is known and that relaxation data can be obtained for sufficiently many sites in the protein.^2,28^ On the computational side, several methods have been developed to investigate rotational diffusion with MD simulations, typically by fitting diffusion models to rotational correlation functions extracted from the trajectories. Early methods used hundreds to thousands of correlation functions of unit vectors that rotate with the molecule.^27,29,30^ More recent approaches utilize quaternions to extract the rotational correlations directly from the orientation of the molecule,^1,31^ which significantly reduces the number of correlation functions. Both methods are capable of extracting the anisotropic diffusion coefficients as well as the principal axes from the correlation functions. Alternatively, the diffusion coefficients can be fitted to the mean angular displacement around the inertia axes, assuming that the principal axes of the diffusion tensor and the inertia tensor coincide.^32^ However, as Wong and Case emphasized,^27^ although it is always possible to fit an anisotropic diffusion model, this does not imply that the rotational dynamics are indeed diffusive or that the fitted model is statistically supported by the data.

Therefore, in this work, we extend the quaternion-based method^1,31^ by introducing a time-dependent analysis of the diffusion coefficients and principal axes from MD simulations. The analysis indicates whether the rotational dynamics observed in MD simulations follow (on average) the motion of a free Brownian rigid rotor as required by Favro’s diffusion theory.^26^ In addition, the time-dependent analysis aids in selecting a suitable diffusion model, anisotropic, semi-isotropic, or isotropic, and a correlation time interval in which the rotational dynamics are Brownian and statistically converged. This time interval is subsequently used to fit a single diffusion tensor using a combined global and local optimization procedure. Finally, we introduce intuitive and interpretable uncertainty metrics for both the diffusion coefficients and the principal axes. We showcase the method by applying it to extended multi-microsecond time scale atomistic MD simulations of the small globular protein ubiquitin in aqueous solution.

## II. THEORY

### A. Extracting rotational correlation functions from MD simulations

Figure 1 illustrates the protocol for extracting rotational correlation functions from MD simulations.^1,31^ First, center-of-mass translational motion is removed from the MD trajectory to isolate rotational movement. Then, all trajectory frames are rotated such that they are optimally aligned to a common reference structure. The corresponding rotation matrices describe the molecule’s orientation and they are converted to unit quaternions

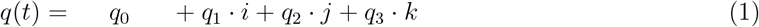

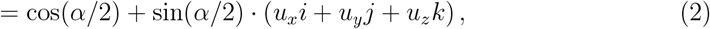

where *i, j*, and *k* are complex numbers, *α* is the rotation angle, and 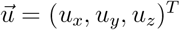 defines the rotation axis. The molecule’s reorientation (or rotation) from time *t* to time *t* + *τ* is given by the quaternion product

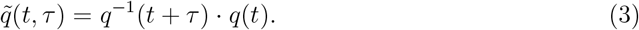

**FIG. 1.**
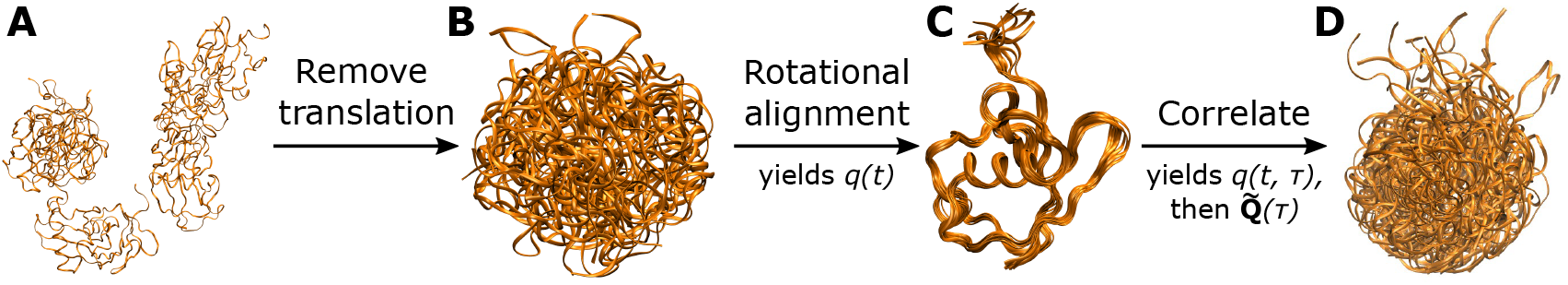
Protocol for extracting rotational correlation functions from MD simulations, illustrated using 20 trajectory frames from simulations of ubiquitin. The alignment step (from **B** to **C**) corre-sponds to a coordinate transform to internal *body* coordinates. **D** visualizes the rotational spreadof the aligned ensemble in **C** within *τ* = 5 ns. 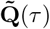 contains the desired correlation functions.

A transformation to internal *body* coordinates yields

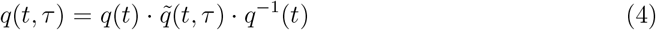

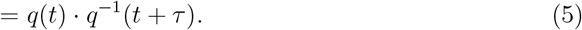

Eq. 5 differs from Eq. 3 because quaternion multiplication is not commutative. The *body* coordinate system rotates with the molecule, so all frames t are aligned (Figure 1C). The reorientations *q*(*t, τ*) in Eq. 5 describe the rotational spread from the aligned ensemble within time *τ* (Figure 1D).

The complex components *q*_1_, *q*_2_, and *q*_3_ of *q*(*t, τ*) form the quaternion covariance matrix

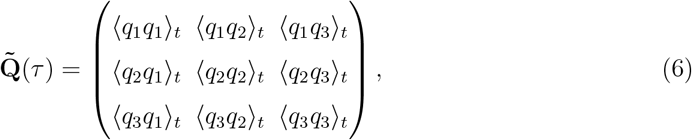

where ⟨…⟩_*t*_ denotes averaging over *t* and *τ* is the correlation time. The matrix corresponds to Eq. (14) in the work by Chen *et al*.^1^ and Eq. (10) in the work by Linke *et al*.^31^ The matrix is symmetric and contains six rotational correlation functions. The tilde symbol (~) denotes that the functions are in *body coordinates*.

Similarly, a matrix of variances defined as

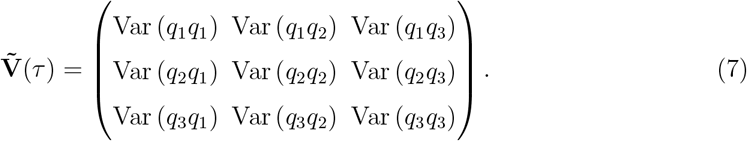

The procedure outlined here for extracting rotational correlation functions loosely follows the work of Chen *et al*.^1^ The procedure by Linke *et al*. is essentially identical, but the conversion from rotational matrices to quaternions happens at a later stage.^31^ An advantage of an early conversion to quaternions, as adopted here, is that quaternions are numerically more stable and computationally more efficient than rotation matrices.

### B. Theory of the rotational Brownian motion of a rigid body

Favro developed a comprehensive mathematical framework describing the anisotropic Brownian rotational motion of a rigid body.^26^ The (unknown) principal axes of the diffusion tensor constitute an orthonormal *principal coordinate system* (**PCS**). In the limit of infinite sampling, the quaternion covariance matrix is diagonal in the **PCS**; in other words, the principal axes are eigenvectors of the quaternion covariance matrix 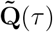. The corresponding matrix-eigenvalue equation

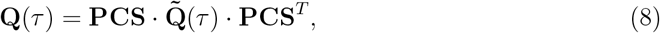

describes the rotation of 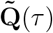 from *body* to *principal* coordinates. The principal axes are the row vectors of **PCS**. The corresponding eigenvalue functions **Q**_*ii*_(*τ*) depend parametrically on the rotational diffusion coefficients,^26,31^

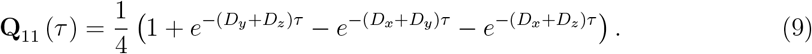

**Q**_22_ and **Q**_33_ follow analogously by permuting *x, y*, and *z* (**Q**_12_, **Q**_13_, and **Q**_23_ are zero since **Q** is diagonal). Favro’s model includes equations to express also the relation between the diffusion coefficients and the variances **V**_*ij*_(*τ*) in *principal* coordinates.^26,31^

### C. Time-dependent principal axes, principal angles, and diffusion coefficients

There is no single **PCS** that solves Eq. (8) for all correlation times *τ*, because the averages extracted from MD simulations suffer from noise due to finite sampling. This problem may be solved by treating both the principal axes and the diffusion coefficients as functions of correlation time *τ*. Then, Eq. (8) can be solved separately for each *τ* to yield the *time-dependent principal axes*, **PCS**(*τ*), which are vector functions. To simplify plotting and analysis, we define the scalar *principal angles*

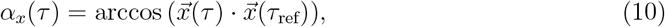

where 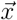 is the time-dependent *x*-axis and *τ*_ref_ is a fixed reference time. *α*_*y*_ and *α*_*z*_ are defined analogously using the *y*-axis and *z*-axis, respectively.

The time-dependent eigenvalues **Q**_*ii*_(*τ*) are used to compute the *time-dependent diffusion coefficients* as

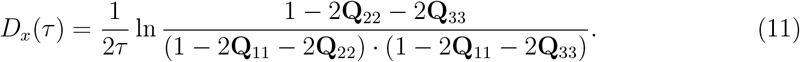

Again, *D*_*y*_(*τ*) and *D*_*z*_(*τ*) follow analogously by permuting 11, 22, and 33. Eqs. (9) and (11) are equivalent when taking all **Q**_*ii*_(*τ*) and *D*_*i*_(*τ*) into account.

### D. Fitting principal axes and diffusion coefficients

For some applications, such as comparing to experimental values, it is necessary to find a single set of diffusion coefficients and principal axes which best characterizes the overall rotational dynamics. Such a set can be obtained using a fitting procedure.^31^

To find the optimal parameters, we minimize the sum of squared residuals,

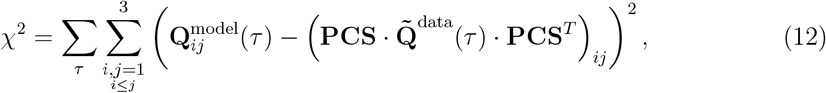

where 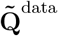 is the quaternion covariance matrix extracted from MD simulations, rotated into the unknown *principal coordinate system* using **PCS. Q**^model^ is the diagonal model covariance matrix, which is computed in each minimization step from the current diffusion coefficients using Eq. (9). To avoid double counting of the off-diagonal matrix elements, the summation index is restricted to *i* ≤ *j*.

The *χ*^2^ function has six degrees of freedom (DoFs), the three diffusion coefficients embedded in the model covariance matrix **Q**^model^, and three rotational DoFs describing the orientation of **PCS**. Optimizing all six DoFs independently corresponds to fitting the anisotropic diffusion model. A semi-isotropic of isotropic diffusion model can be fit by enforcing two or all three diffusion coefficients, respectively, to be equal. Thereby, the number of DoFs is reduced to five or four, respectively, and (some of) the obtained principal axes have no physical meaning.

The residuals function in Eq. (12) differs from the function used by Linke *et al*.,^31^ which in our notation is

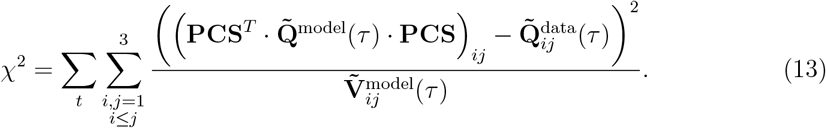

First, the residuals in Eq. (13) are computed in *body* coordinates, so the residuals depend on the orientation of the reference structure, which, consequently, also applies to the diffusion parameters. In contrast, the residuals in Eq. (12) are computed in *principal* coordinates, which are intrinsically independent of the reference orientation. Second, the residuals in Eq. (13) are weighted using the variances from Eq. (7). We decided against this weighting because the majority of the weights are accumulated for very few initial data points close to *τ* = 0, since the variances are initially close to zero and the weights are correspondingly large.

Both residuals functions in Eqs. (12) and (13) are nonlinear, indicating that multiple minima may exist. Hence, Linke *et al*. developed a simulated annealing algorithm that searches for the global minimum.^31^ The algorithm terminates close to but not precisely at a local minimum in a stochastic way. We propose a local optimization algorithm that deterministically identifies the next local minimum given an initial guess. The two algorithms can be used separately or together, performing an initial global parameter search before finishing with a local optimization. We discuss alternative optimization strategies in Sec. IV C.

Chen *et al*. proposed an alternative method for fitting an isotropic diffusion model to the mean rotation angle cosine, which is directly accessible from the trace of the quaternion covariance matrix.^1^ However, concerning the anisotropic model, Eq. (16) from Chen *et al*.^1^ is not equivalent to Eq. (6.2) in Favro’s work^26^ (Eq. (9) above). Here, we stick to fitting diffusion models directly to the components of the quaternion covariance matrix instead of its trace, which provides a consistent framework for fitting anisotropic, semi-isotropic, or isotropic diffusion models.

### E. Finite size and viscosity corrections

Rotational dynamics in MD simulations suffer from two error sources that can be analytically corrected. First, rotational diffusion is slowed down because the finite size of the periodic simulation box creates an artificial drag. The additive correction is^33^

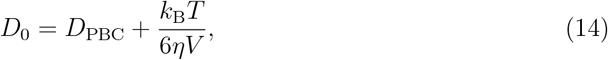

where *k*_B_ is the Boltzmann constant, *T* the temperature, *η* the viscosity of the solvent, and *V* the volume of the simulation box. *D*_0_ and *D*_PBC_ are the corrected and uncorrected diffusion constants, respectively.

The Stokes-Einstein-Debye relation

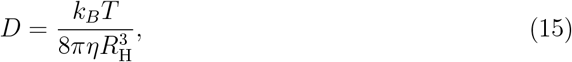

where *R*_H_ is the hydrodynamic radius of the solute, allows for the definition of a further correction:

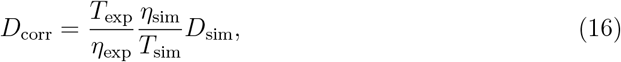

where *T*_exp_, *T*_sim_, *η*_exp_, and *η*_sim_ are the experimental and simulated temperatures and viscosities, respectively.Eq. (14) can be used to scale the diffusion coefficients to a different temperature, e.g., for comparison with experimental data. More importantly, the relation accounts for errors in the viscosity of the water model.

In contrast, other errors in the simulation cannot be analytically corrected. For example, erroneous sampling of the conformational space due to insufficient sampling or force field errors may affect the rotational dynamics. In addition, unbalanced water-protein and water-water interactions can lead to water molecules attaching overly tight (or loose) to the protein,^34^ which might artificially slow down (or speed up) the rotational dynamics.

### F. Brownian dynamics simulations and uncertainty metrics

The statistical uncertainty of the obtained rotational diffusion coefficients and principal axes can be estimated using rotational Brownian dynamics (BD) simulations.^31^ BD simulations are several orders of magnitude more efficient than MD simulations and thus better suited for determining the uncertainty. A BD simulation uses a reference set of diffusion coefficients and principal axes to generate a stochastic trajectory of orientations *q*(*t*), intrinsically obeying the laws of Brownian rotational motion. The BD trajectory can be analyzed analogously to the trajectory of orientations obtained from MD to yield a new set of diffusion coefficients and principal axes. In general, these new results differ from the reference coefficients and axes due to the stochastic nature of Brownian motion.

The statistical uncertainty of the fitted diffusion parameters due to finite sampling of the Brownian process can be assessed by running many BD simulations using the MD-derived parameters as the reference. The simulations yield a distribution of diffusion parameters spread around the reference parameters, following an unknown multi-dimensional joint probability density. We approximate the probability density as disjoint, which allows one to examine the uncertainties of individual DoFs. The uncertainty of the three diffusion coefficients is then simply their standard deviation. Linke *et al*. used the Frobenius norm to compare diffusion tensors,^31^ which considers correlations between the diffusion coefficients and therefore it cannot distinguish the uncertainties of the individual coefficients.

The uncertainties of the three remaining DoFs indicate how precisely the orientation of the principal axes is known. Again, we treat the three principal axes individually. Thereby, the number of orientational DoFs is effectively increased from three to six, because the orthogonality constraints of the axes are lost. Correspondingly, six uncertainties are obtained, two for each axis. The uncertainties of the *x*-axis are obtained by projecting each BD *x*-axis into the *xy*- and *zx*-planes. The projection is performed in *principal coordinates* defined by the reference **PCS**. Then, the angles between the projected vectors and the reference *x*-axis are computed. The standard deviation of these angles are the *directional* uncertainties of the x-axis in the *xy*- and *zx*-planes. For simplicity, we denote these as the *x-xy* and *x-zx* uncertainties, respectively. The directional uncertainties of the *y*- and the *z*-axis are defined analogously.

## III. COMPUTATIONAL METHODS

### A. Molecular dynamics (MD) simulations

The MD simulation system was prepared using GROMACS 2023.4^35^ and ambertools 23.6^36^. The initial atomic coordinates of ubiquitin were obtained from the protein data bank (PDB ID: 1UBQ^37^). Hydrogen atoms were added before placing the protein structure in a rhombic dodecahedron box with 1.2 nm minimum distance from the protein to the box edges. The system was solvated with 7043 water molecules and 21 sodium chloride ion pairs were added to achieve a physiological ion concentration of approximately 150 mM.

The Amber ff19SB force field^38^ was used for the protein, combined with the OPC3 water model^39^ and Li and Merz ion parameters^40^. An in-house python script was used to convert the residue-specific CMAP parameters from Amber to GROMACS topology format, to ensure the proper implementation of the residue-specific CMAPs defined in the ff19SB force field.

MD simulations were performed under periodic boundary conditions with GROMACS 2025.1^35^. The system underwent three stages of energy minimization using steepest descent: first with harmonic position restraints (force constants 1000 kJ/mol/nm^2^) on all protein heavy atoms, then on backbone heavy atoms only, and finally without restraints.

Ten replicate simulations were initiated from the minimized structure with velocities drawn from a Maxwell-Boltzmann distribution at 300 K. The systems were equilibrated for 150 ps, gradually releasing position restraints every 50 ps according to the minimization protocol. Ten production runs of one microsecond per replica followed.

The temperature was maintained at 300 K using velocity rescaling^41^ (0.1 ps coupling time constant). Coupling to a pressure bath (1 bar) was switched on after 50 ps equilibration using the Parrinello-Rahman barostat^42,43^ (5 ps coupling time constant, 4.5 × 10^−5^ bar^−1^ compressibility). The van der Waals forces were switched to zero between 1.0 nm and 1.2 nm. Long-range electrostatics were handled with the smooth particle mesh Ewald method^44–46^ (0.12 nm grid spacing). Center-of-mass translation was removed every 100 steps. In the protein, bonds to hydrogen atoms were constrained with LINCS^47^ using a sixth-order expansion of the constraint coupling matrix and two iterations to correct for rotational bond lengthening. SETTLE^48^ was used to constrain the internal degrees of freedom of water molecules. The equations of motion were integrated with 2 fs time steps using the leap frog integrator.

### B. Analyzing rotational diffusion from MD

The rotational diffusion of ubiquitin was analyzed using our python package *rotationaldiffusion* (https://github.com/MolSimGroup/rotationaldiffusion). The analyzes were restricted to the backbone atoms *C, C*_*α*_, and *N*. If not indicated otherwise, 10^4^ trajectory frames with an equidistant spacing of 100 ps were used per replicate simulation for the analysis. Center-of-mass translation was removed before aligning the frames to the energy minimized structure using Theobald’s QCP method^49,50^ implemented in MDAnalysis.^51^ The obtained rotation matrices were converted to quaternions *q*(*t*) using a custom implementation of Bar-Itzhack’s algorithm.^52^ The quaternion covariance matrix (Eq. 6) was computed up to 50 ns and averaged over all ten replicates. The time-dependent diffusion coefficients and principal angles (*τ*_ref._ = 1 ns) were used to choose an appropriate fit window from 100 ps to 10 ns (see below).

Anisotropic, semi-isotropic, and isotropic diffusion models were fitted to the rotational correlation functions between 100 ps and 10 ns by minimizing Eq. 12, as described in Sec. III C. The finite size and viscosity corrections were applied to all obtained diffusion coefficients. 1000 sets of BD simulations were performed with *pydiffusion*^31^ using the fitted, uncorrected anisotropic diffusion model as a reference. Each BD simulation set consisted of ten independent replicates sampled for 1 µs each, identical to the MD protocol. Rotational correlation functions were extracted from the BD simulations, fitted analogous to the MD data, and the finite size and viscosity corrections were applied. The uncertainties of the MD diffusion coefficients and principal axes were obtained from the distributions of BD fitting parameters (standard deviations).

To investigate the influence of the residuals function and the fit window, the fitting procedure was repeated using the anisotropic diffusion model, both residuals functions (Eq. (12) and Eq. (13)), separately, and using the correlation functions between 100 ps and 10 ns with a 100 ps spacing, between 100 ps and 50 ns with a 100 ps spacing, or between 1 ps and 10 ns with a 1 ps spacing. In addition, to investigate the influence of the reference structure used for aligning the trajectory frames, the analyzes were repeated three times using different structures as the reference (instead of the energy-minimized structure). The first alternative reference structure was the average structure, which was determined iteratively by aligning the trajectory frames before averaging over all atomic positions. The energy-minimized structure served as the initial reference for the alignment. After three iterations, the averaging scheme was converged with a backbone RMSD change of less than 10^−3^ Å.

The second structure was the trajectory frame with the lowest RMSD with respect to the average structure. For the third structure a matrix was computed containing the RMSDs of all pairs of trajectory frames after alignment. The analysis was restricted to 1000 frames by using every 10th frame, because the computational effort scales quadratically. The RMSDs were averaged over all rows, yielding the mean RMSD of each frame with respect to all other frames. We chose the frame with lowest mean RMSD as the third reference structure. The anisotropic diffusion model was fitted to the rotational correlation functions obtained using each of the different reference structures. The fit interval was kept between 100 ps and 10 ns and Eq. (12) was minimized during the optimization. Again, the finite size and viscosity corrections were applied to all obtained diffusion coefficients.

### C. Optimization of diffusion parameters

A two-step optimization procedure was used to fit diffusion models to rotational correlation functions extracted from MD simulations. The target function to be minimized was either Eq. (12), or Eq. (13). The initial guess for the optimization parameters was obtained from the time-dependent solution as the last data point at which all rotational correlation functions were still below 0.1.

In the first optimization step, the parameter space was explored globally using a simulated annealing algorithm, similar to the one used by Linke *et al*.^31^ In each step of the optimization, either the principal axes or the diffusion coefficients were updated based on a Metropolis criterium. The algorithm switched between updating the axes or the coefficients every 20 steps. Initially, the principal axes were allowed to rotate by at most 90°per step and the diffusion coefficients could change by up to 50 % of their current value. These numbers were linearly reduced to 1°and 5 %, respectively, within the first 2500 out of 10000 optimization steps. Similarly, the annealing temperature was linearly decreased to reduce the probability of accepting a step uphill the target function. Initially, an increase of 1000 % was accepted with a probability of 50 %, which was lowered to an increase of 1 % within 2500 steps. In total, ten independent annealing runs were performed for 10000 steps each, yielding ten sets of parameters.

In the second step, each set of parameters was further optimized locally using the Byrd-Omojokun trust region SQP method^53^ implemented in scipy. Thereby, the **PCS** was represented as a quaternion, constraining its norm to unity. The diffusion coefficients were represented by their decadic logarithm to match the order of magnitude to the quaternion parameters, which allowed optimizing all parameters at once. Ten sets of locally optimized parameters were obtained, one for each annealing run.

To fit diffusion models to rotational correlation functions extracted from BD simulations, we skipped the simulated annealing step and used the local minimization algorithm to obtain one set of optimized parameters.

## IV. RESULTS AND DISCUSSION

### A. Rotational correlation functions

We conducted ten MD simulations of ubiquitin in explicit solvent, each MD trajectory 1 µs long, employing the Amber ff19SB protein force field^38^ and the OPC3 water model^39^. Utilising our Python package *rotationaldiffusion*, we examined the rotational dynamics of ubiquitin in the MD simulations.

First, the quaternion covariance matrix 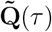 and variance matrix 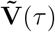 were extracted from the simulations (Fig. 2). The rotational correlation functions in 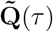 qualitatively follow Eq. (9) although they are in *body* coordinates. The diagonal functions in 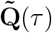 begin at zero, rise, and eventually converge to values near 0.25 in the long correlation time limit. The off-diagonal functions remain close to zero. The variances 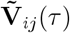 begin at zero, increase, and eventually converge.

**FIG. 2.**
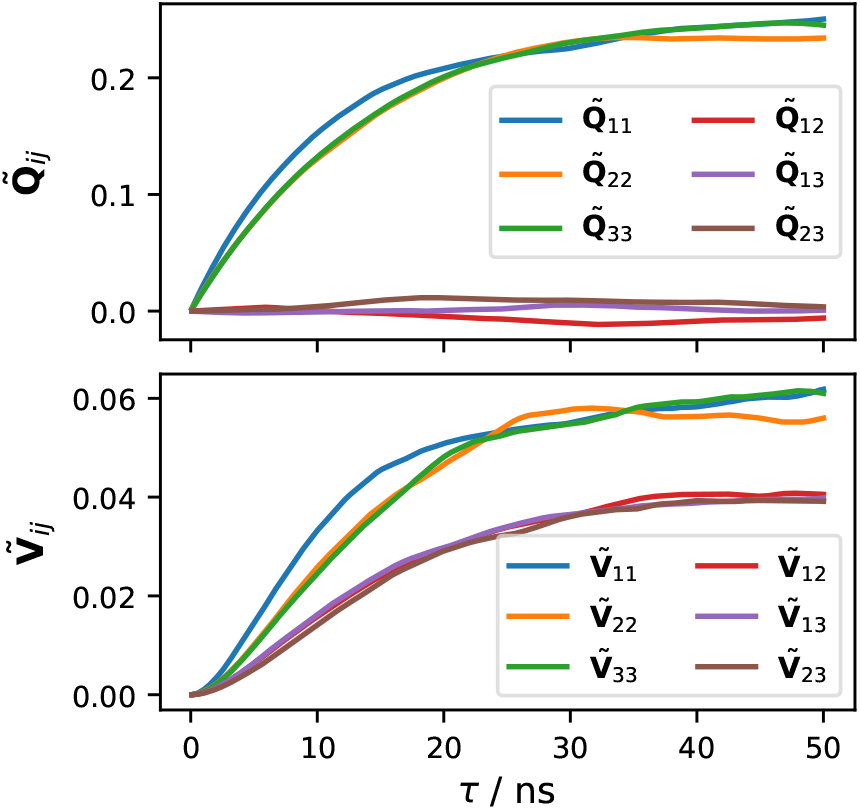
Rotational correlation functions (top) and variances (bottom) extracted from MD simulations of ubiquitin according to Eqs. (6) and (7), respectively.

### B. Time-dependent diffusion coefficients and principal angles

The time-dependent rotational diffusion coefficients and principal angles of ubiquitin (Fig. 3) were computed from the rotational correlation functions in Fig. 2. For comparison, we also performed BD simulations using the fitted anisotropic diffusion model as input (see Sec. IV C) and included the results of the time-dependent analysis in Fig. 3. Beyond 100 ps, the MD-derived diffusion coefficients behave qualitatively identical to the BD coefficients, indicating that Favro’s diffusion theory accurately describes the rotational motions observed in MD. The coefficients are constant up to approximately 10 ns before deteriorating, similar to the BD coefficients. Notably, the deviations start at a correlation time at which the rotational dynamics are still significantly correlated (Fig. 2).

**FIG. 3.**
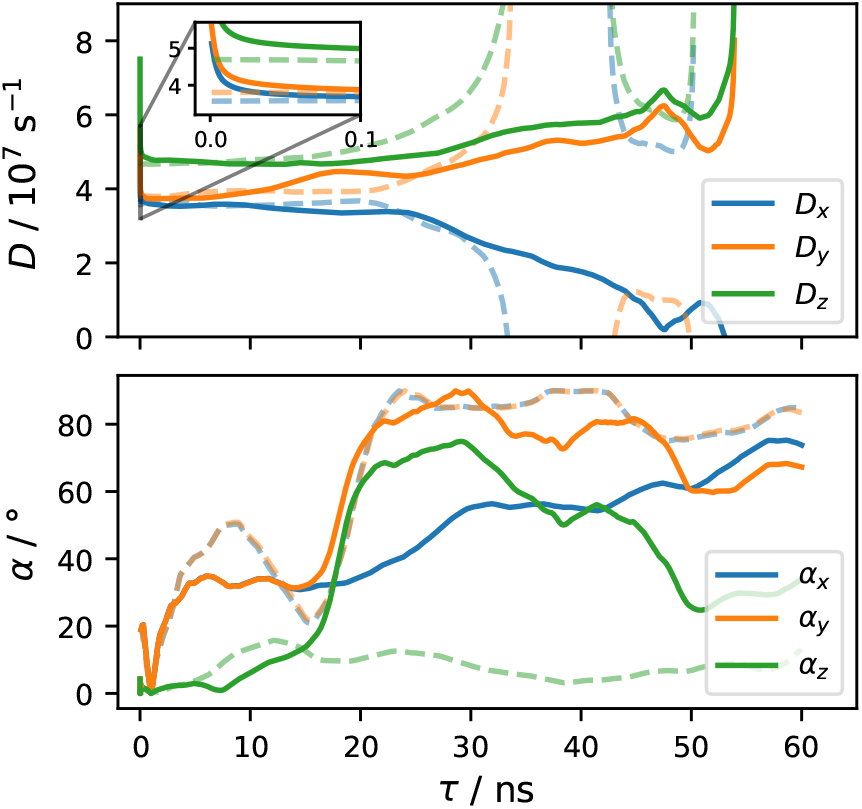
Time-dependent diffusion coefficients (top) and time-dependent principal angles (bottom) extracted from MD simulations (solid lines) or from BD simulations (dashed lines). The BD simulations used the fitted anisotropic diffusion model as input. The principal angles were computed using Eq. (10) with *τ*_ref._ = 1 ns. The spacing between data points is in general 100 ps, but 1 ps for data below 100 ps to resolve the short time region.

At short correlation times below 100 ps, the MD-derived diffusion coefficients exhibit a steep initial drop, contradicting the BD coefficients, which are constant. This finding is in line with the results of Ollila *et al*., who reported non-Brownian rotational diffusion for time scales faster than 120 ps for the TonB C-terminal domain.^32^ Apparently, Favro’s diffusion theory is not sufficient to explain the rotational motions on such short time scales.

Both the diffusion coefficients and principal angles from MD suggest that the rotational dynamics of ubiquitin are close to semi-isotropic. The *D*_*x*_ and *D*_*y*_ coefficients are similar and their corresponding principal angles deviate substantially from zero even below 10 ns, which indicates degenerate diffusion modes.

The deterioration of both the MD- and BD-derived diffusion coefficients beyond 10 ns is a consequence of limited statistics due to finite sampling, which affects the diffusion coefficients in two ways. First, the uncertainty of the rotational correlation functions increases with increasing correlation time, as illustrated by the variances in Fig. 2. The increase in uncertainty is propagated to the diffusion coefficients. Second, the rotational correlation functions 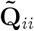 approach 1/4 with increasing correlation time (Eq. 9), causing the argument in the logarithm in Eq. (11) to approach a “zero-by-zero” division. In consequence, minor statistical fluctuations in the rotational correlation functions propagate to large changes in the diffusion coefficients. The coefficients become undefined when the argument becomes negative, as observed in the BD coefficients between 33 and 43 ns (Fig. 3). In summary, the deterioration of the diffusion coefficients confirms that the rotational dynamics decorrelate increasingly and the rotational correlation functions provide no further valuable information at long correlation times.

### C. Fitted diffusion coefficients and principal axes

We fitted all three diffusion models, anisotropic, semi-isotropic, and isotropic, to the rotational correlation functions of ubiquitin. Based on the time-dependent analysis, we limited the fit window to data in the correlation time interval up to 10 ns with a 100 ps spacing, thus excluding the fast dynamics below 100 ps. In addition to the diffusion coefficients, we report the anisotropy^54^ the rhombicity^54^

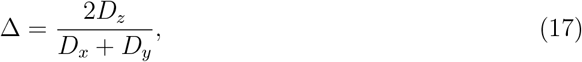

the rhombicity^54^

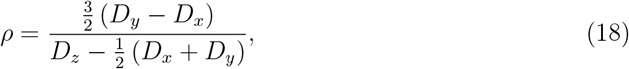

and the isotropic rotational correlation time^54^

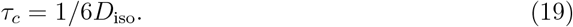

All obtained parameters are collected in Table I, and the principal axes with respect to the reference structure are shown in Figure 4. The *z*-axis passes the termini of the polypeptide chain. The finite box size correction for the diffusion coefficients is 0.40 × 10^7^ s^−1^, using the simulation temperature of 300 K, average box volume of 224.4 nm^3^, and viscosity of OPC3 water at 300 K of 0.77 × 10^−3^ Pa s.^55^ The experimental viscosity is 0.85 × 10^−3^ Pa s^56^ at 300 K, resulting in a multiplicative viscosity correction of 0.906.

**TABLE 1.**
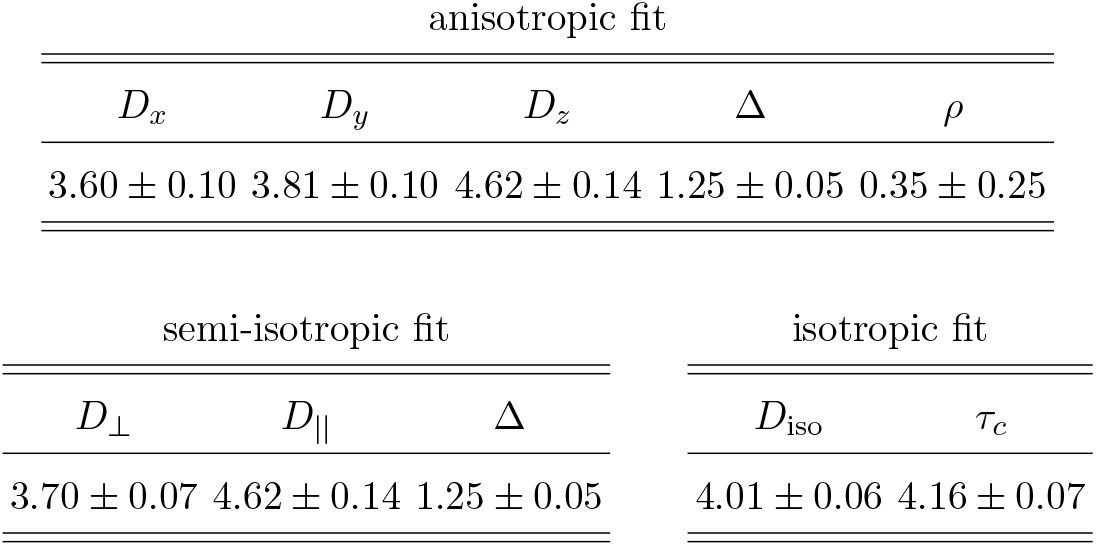
Rotational diffusion parameters of ubiquitin obtained by fitting model functions to the rotational correlation functions between 100 ps and 10 ns by minimizing Eq. (12). Diffusion coefficients are reported in 10^7^ s^−1^, the anisotropy Δ and rhombicity *ρ* are dimensionless, and the isotropic rotational correlation time *τ*_*c*_ = 1*/*6*D*_iso_ is in nanoseconds. The reported diffusion coefficients have been corrected for finite box size and viscosity (Eqs. (14) and (16)). The uncertainties were obtained from 1000 sets of Brownian dynamics simulations.

**FIG. 4.**
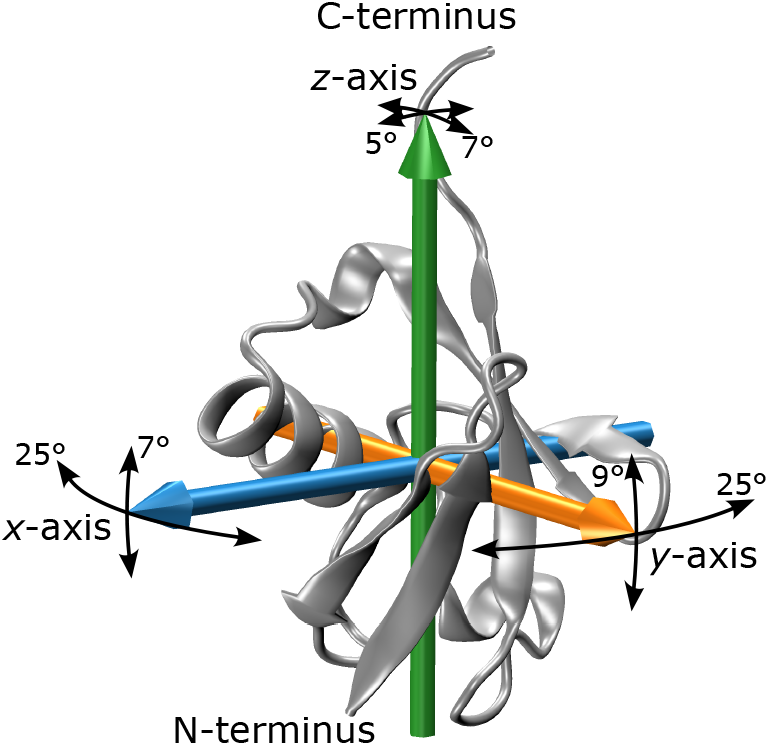
Principal axes of the fitted anisotropic diffusion tensor of ubiquitin. The thin black arrows indicate the directional uncertainties extracted from 1000 BD simulations (arrow lengths are proportional to the corresponding uncertainties).

The uncertainties were obtained from 1000 BD simulations. The same corrections used for the MD coefficients were also applied to the BD diffusion coefficients. The resulting distributions of BD diffusion coefficients and angles of principal axes are depicted in Figure 5. The relative uncertainties of the MD diffusion coefficents are at most 3% (Table II). However, there is a significant overlap of the *D*_*x*_ and *D*_*y*_ coefficients and their corresponding angles in the *xy*-plane. In fact, there is a 9% probability that the actual *x*-axis is closer to the obtained *y*-axis and vice versa, due to the statistical noise. In contrast, the angular distributions in the *yz*- and *zx*-planes are sharp, indicating that diffusion around the *z*-axis is significantly different from diffusion around the *x*- and *y*-axes. Thus, when computing, for example, NMR relaxation parameters from MD simulations, it may be advisable to use a statistically more robust albeit more approximate semi-isotropic diffusion model instead of the fully anisotropic one.^23^

**TABLE 2.** Rotational diffusion parameters of ubiquitin obtained by fitting an anisotropic diffusion model to the rotational correlation functions, varying the residuals function *χ*^2^, the maximum correlation time *τ*_max_, and the data spacing Δ*τ*. Δ*τ* and *τ*_max_ are reported in nanoseconds and the diffusion coefficients in 10^7^ s^−1^.

| $\chi^2$ | $\Delta\tau$ | $\tau_{\text{max}}$ | $D_x$ | $D_y$ | $D_z$ |
| --- | --- | --- | --- | --- | --- |
| Eq. (12) | $10^{-1}$ | 10 | 3.60 | 3.81 | 4.62 |
| Eq. (12) | $10^{-3}$ | 10 | 3.60 | 3.81 | 4.62 |
| Eq. (12) | $10^{-1}$ | 50 | 3.45 | 4.24 | 4.51 |
| Eq. (13) | $10^{-1}$ | 10 | 3.61 | 3.77 | 4.66 |
| Eq. (13) | $10^{-3}$ | 10 | 3.61 | 3.78 | 4.66 |
| Eq. (13) | $10^{-1}$ | 50 | 3.59 | 4.04 | 4.53 |

**FIG. 5.**
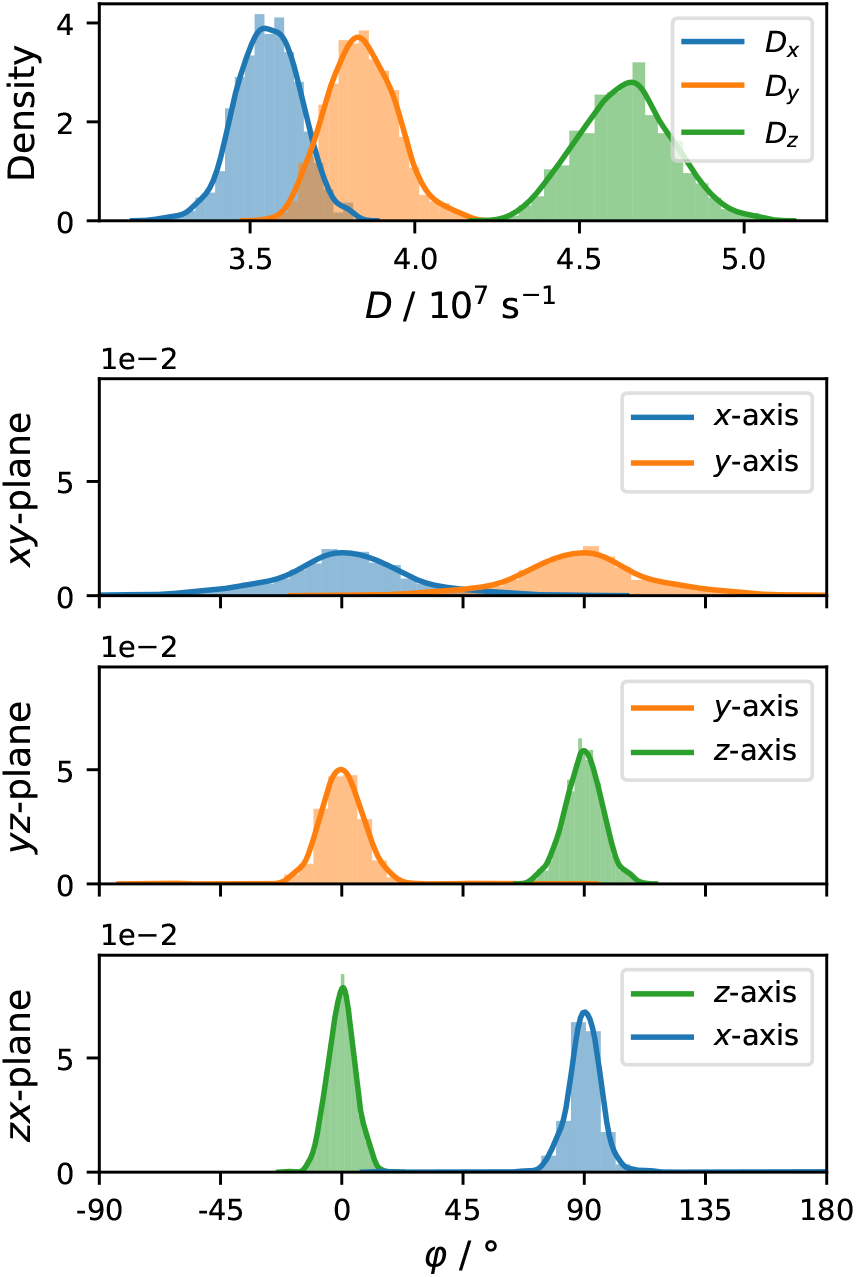
Distributions of diffusion coefficients (top plot) and angles of principal axes after projection into the indicated coordinate planes (bottom three plots) obtained from anisotropic fits of 1000 BD simulations.

The obtained isotropic diffusion coefficient of (4.01±0.06)×10^7^ s^−1^ agrees well with the range of values (in 10^7^ s^−1^) from NMR experiments of 4.01,^57,58^ 4.05,^2^ and 4.14.^59^ The anisotropy of 1.25 ± 0.05 is slightly higher than the experimental values of 1.15,^57^ 1.17,^2^ and 1.18.^58^ This small difference may be within the limits of the statistical uncertainties.

### D. Influence of the residuals function, fit window, and reference structure

To investigate the impact of the residuals function (Eq. (12) or Eq. (13)) and the fit window on the anisotropic diffusion coefficients, we repeated the fitting procedure with different input parameters and report the obtained coefficients in Table II. The two residuals functions yield almost identical diffusion coefficients when keeping the fit window between 100 ps and 10 ns. The coefficients differ by less than 1.1%. The two functions differ mainly in their weighting scheme, which has no effect because the fit window was chosen such that the time-dependent diffusion coefficients are almost constant.

Including the very fast dynamics below 100 ps in the fitting procedure (by using a data spacing of 1 ps) does not alter the obtained diffusion coefficients, regardless of the residuals function. Conversely, including noisy parts of the correlation functions by extending the fit window to 50 ns yields diffusion coefficients that significantly differ from the ones obtained before. In addition, the coefficients now depend on the residuals function. The coefficients obtained using Eq. (12) suggest that ubiquitin is more oblate than prolate (Table II), contradicting experimental results.^2^ The coefficients obtained using Eq. (13) indicate anisotropic rotational diffusion with *D*_*y*_ being significantly larger than *D*_*x*_, which is again contradicting the experiments. These findings underline the relevance of limiting the fitting procedure to a correlation time window in which the rotational dynamics are still sufficiently correlated and statistically converged.

Finally, we repeated the fitting procedure using three alternative reference structures. In addition to the minimized structure, we considered the average structure, the trajectory frame with the lowest RMSD with respect to the average structure, and the trajectory frame with the lowest average RMSD with respect to all other frames. The structures barely differ, as shown in Fig. 6. It is therefore not surprising that the results of the rotational diffusion analysis are almost identical. The anisotropic diffusion coefficients agree in at least three significant digits, and the principal axes are within 0.4°despite the large uncertainties. The remaining differences are negligible considering the significantly larger uncertainties.

**FIG. 6.**
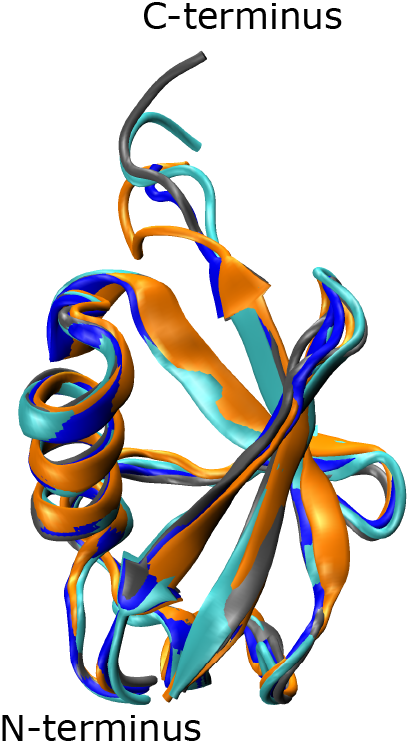
Superimposed reference structures used for the analysis, including the minimized structure (grey), the average structure (blue), the trajectory frame with lowest RMSD with respect to the average structure (cyan), and the frame with lowest mean pairwise RMSD (orange).

Nevertheless, choosing a suitable reference structure may become more important, for example, when investigating flexible molecules, where the reference structure types used here are expected to exhibit more variation. In that case, the trajectory frame closest to the average structure is superior to the others from a theoretical point of view. It is the only structure considered here that incorporates information of the whole structural ensemble (because it is derived from the average structure), is physically meaningful, and easy to compute.

### E. Comparison of the optimization algorithms

We employed a two-step optimization procedure to minimize the residuals functions, Eq. (12) or Eq. (13). The first algorithm is a simulated annealing algorithm, similar to the one used by Linke *et al*.^31^ The algorithm enables jumps between local minima in the initial phase of the optimization, which is important, because the residuals functions are six-dimensional and nonlinear in their parameters. Hence, in theory, several local minima may exist. Afterwards, the algorithm smoothly transitions to descending into a local minimum. The algorithm explores the parameter space stochastically and may terminate close to, but not precisely at, the exact minimum.

Therefore, we used a local optimization algorithm after the simulated annealing. The algorithm deterministically locates the next local minimum given an initial guess. We used the two algorithms together for fitting the diffusion models to the MD data, by first exploring the full parameter space in several runs of the simulated annealing algorithm before descending into the local minimum with the local optimization algorithm. In all cases, the results after local optimization were identical up to numerical accuracy, verifying that all runs of the annealing algorithm ended up in the same minimum, as illustrated in Fig. 7. In fact, conducting the local minimization without prior simulated annealing yielded the same minimum. Apparently, the residuals functions have a single minimum in the present case and the simulated annealing step is not required.

**FIG. 7.**
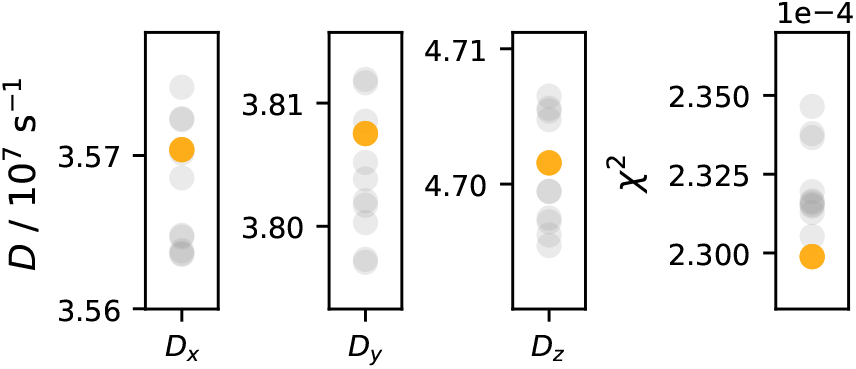
Optimized diffusion coefficients (left three plots) after ten independent runs of simulated annealing (grey) or a subsequent local minimization (orange). Eq. (13) was minimized during the optimization; the final values are presented in the plot on the right.

In general, the combined optimization procedure is superior to a single local optimization, because it works also if the residuals function has multiple minima. Nonetheless, skipping the annealing phase is preferential if computational efficiency is of interest, for example, for fitting the data from very many BD simulations to estimate uncertainties.

Based on the MD results, we assumed that the data from BD simulations may be fitted using only the local optimization algorithm. To further verify this approach, we used 10 of the 1000 BD simulations and repeated the anisotropic fitting 100 times each with random initial guesses. The initial principal axes were chosen as a random rotational matrix. We used the fitted diffusion coefficients as input to target the correct order of magnitude, but varied each coefficient by applying random factors between 0.1 and 10. All fits of the same BD simulation yielded numerically identical results (up to at least five significant digits). We conclude that the local optimization suffices, at least in the present case.

## V. SUMMARY AND CONCLUDING REMARKS

In this work, we present a time-dependent analysis of the rotational diffusion of molecules from MD simulations. The approach is illustrated using atomistic MD simulations of the small globular protein ubiquitin in explicit water. It provides a systematic way to investigate whether the rotational dynamics are Brownian (as required by Favro’s theory) and to choosean appropriate diffusion model (isotropic, semi-isotropic, or anisotropic) that is supported by the data given the statistical noise due to finite sampling.

With the 10 microsecond time scale MD simulations carried out in the present work, the time-dependent analysis revealed that the rotational correlation functions of ubiquitin are converged up to correlation times of 10 ns, which is approximately twice the rotational correlation time of ubiquitin in water at room temperature. Thus, the diffusion models were fitted to the data within this suitable time interval. Oversampling the rotational correlation time by about three orders of magnitude, as done in the present case, is exhaustive and exceeds the previously suggested oversampling of two orders of magnitude,^60^ reducing the statistical uncertainties to below 3%. These small errors support the choice of a semiisotropic model with one fast and two slow diffusion modes to describe the rotational diffusion of ubiquitin. The diffusion coefficients obtained from the MD simulations are in quantitative agreement with experimental data from NMR spectroscopy.

The presented approach is robust and it can be useful to analyze the rotational dynamics of different (bio-)molecules under various conditions. In particular, it will be interesting to investigate more flexible biomolecules or non-homogeneous environments, where Favro’s model assumption of a rigid body moving in an isotropic environment is violated. The presented approach is applicable nevertheless, yielding diffusion models of hypothetical rigid bodies, which are averaged over the relevant conformations or environments. In this context, the time-dependent approach is expected to aid in identifying convergence issues, drifts, and non-Brownian dynamics, which are not evident from the correlation functions alone.

## ACKNOWLEDGMENTS

This work was supported by Deutsche Forschungsgemeinschaft (DFG) under Germany’s Excellence Strategy - EXC 2033 - 390677874 - RESOLV.

## AUTHOR DECLARATIONS

The authors have no conflicts to disclose.

## DATA AVAILABILITY

The software package *rotationaldiffusion* developed as part of this work is freely available at https://github.com/MolSimGroup/rotationaldiffusion. The data that support the findings of this study are available from the corresponding author upon reasonable request.

